# Biochemical characterization of extended-spectrum β-lactamases from *Akkermansia* genus

**DOI:** 10.1101/2024.06.10.598323

**Authors:** Jiafu Lin, Tiantian Wang, Yaliang Zhou, Jingzhou Sha, Xueke Chen, Wenjie Wang, Chuan Zhang, Feng Xie, Yiwen Chu, Xinrong Wang, Dan Luo, Tao Song

## Abstract

*Akkermansia muciniphila*, a member of the *Verrucomicrobiota* phylum, is recognized as a key gut microbe and has emerged as a potential next-generation probiotic. Assessment of antibiotic resistance in probiotics is a prerequisite for their therapeutic application, while very few is known in *Akkermansia* species. Firstly, we screened eight representative class A β-lactamases (36.90%-41.30% identity with known β-lactamases) from the *Akkermansia* species, which could increase the minimum inhibitory concentration (MIC) of *Escherichia coli* toβ-lactams. Secondly, fourβ-lactamases were purified and identified as extended-spectrum β-lactamase because they exhibited hydrolase activity against 19 β-lactam antibiotics from penicillin, cephalosporins, and monobactam classes. Based on sequence alignment, three-dimensional structure, and binding pocket information, we hypothesized and validated that serine at 51 position was catalytic amino acid. Thirdly, the genomic context analysis revealed the absence of mobile genetic elements or other antibiotic resistance genes surrounding β-lactamase genes, suggesting that the β-lactamases from *Akkermansia* species may not be transferable. The finding and biochemical characterization of β-lactamase from *Akkermansia* species provide a foundational basis for the safety evaluation of *Akkermansia* species as probiotics.

## 1 Introduction

*Akkermansia muciniphila*, a pivotal species in the human gut microbiome, has garnered attention as an emerging next-generation probiotic(1, 2). First identified in 2004, *Akkermansia muciniphila* belongs to the *Verrucomicrobia* phylum and is notable for its ability to metabolize the mucus layer in the gut(3). This unique function is critical for maintaining gut integrity and health, impacting human health and disease(4, 5). Its interaction with the host’s immune system is a key factor in its influence on health. Disruptions in its population have been linked to various diseases, including obesity, type 2 diabetes, and inflammatory bowel diseases(6–8). These associations are attributed to its regulatory effects on both metabolism and immune responses, highlighting its importance in gut and human health. Numerous studies underscore the potential of *Akkermansia muciniphila* as a next-generation probiotic, however, concerns about its safety, including issues like antibiotic resistance and biofilm formation, remain(9).

The European Food Safety Authority emphasizes the importance of assessing antibiotic resistance in probiotics before their industrial use(10). Gut probiotics, such as *Akkermansia muciniphila*, are highly regarded for their health benefits, yet they pose challenges in terms of antibiotic resistance(9). These beneficial bacteria, which are essential for gut health, may carry genes conferring resistance to antibiotics, including *β*-lactams, tetracycline, macrolides, and aminoglycosides(9, 11, 12). Increasing evidence suggests that these beneficial bacteria can serve as reservoirs for such genes, potentially enabling their transfer to pathogenic bacteria within the gut. This gene transfer could reduce the effectiveness of antibiotic treatments, extending treatment times and cycles, and in some cases, resulting in treatment failure(13, 14). Consequently, despite the substantial health benefits conferred by gut probiotics, their capacity to carry and transmit antibiotic-resistant genes constitutes a significant challenge. This underscores the necessity for comprehensive resistance evaluation prior to their application.

Many research has been conducted on the antibiotic resistance and resistant genes of *Akkermansia muciniphila* (9, 12, 15, 16). Among them, *β*-lactam resistance of AKK is the most studied characteristics due to the extensive use and long history of *β*-lactam antibiotics(17). For instance, *Akkermansia* sp. DSM 33459 has a MIC greater than 32 μg/mL for ampicillin and >16 μg/mL for penicillin, while *A. muciniphila* DSM 22959 is sensitive to ampicillin and imipenem, but resistant to penicillin G(18) (15) (19). Meanwhile, genomic sequencing of *Akkermansia muciniphila* indicated the presence of potential β-lactamase(18). Yet, there are no studies on the biochemical characteristics, enzymatic properties, or catalytic mechanisms of β-lactamase from *Akkermansia muciniphila*.

Assessing antibiotic resistance is a crucial prerequisite for the application of probiotics. Although some *Akkermansia muciniphila* were reported to have β-lactam resistance and β-lactamases were bioinformatically found in their genomes, a systematic and biochemical characterization of their β-lactamases is lacking. In our research, we conducted the first systematic study of β-lactamase from *Akkermansia* sp., screened and biochemically studied 8 representative β -lactamase from *Akkermansia* sp. The genomic context of β-lactamases from *Akkermansia* species was also investigated to infer the transferability of resistance genes. This research pioneers the study of β-lactamases from *Akkermansia*, laying a foundation for safety evaluation.

## 2. Results

### 2.1 Systematically screen of β-lactamases from *Akkermansia* genus

To identify potential β-lactamases in *Akkermansia* species, we first constructed a genome database consisting of 2,659 *Akkermansia* genomes, including 60.10% *Akkermansia muciniphila*, 39.26% *Akkermansia* sp., 0.49% *Akkermansia massiliensis*, 0.11% *Akkermansia glycaniphila*, and 0.038% *Akkermansia biwaensis* (supplementary material 1). Of these genomes, 81.19% were from pure-culture strains and 4.76% were from metagenomes. In total, 1,601,551 protein sequences were extracted and compared with those in the β-lactamase Database (BLDB), which includes 831 from Class A, 637 from Class B, 156 from Class C, and 380 from Class D. Within our dataset, 335 proteins exhibited sequence identity to Class A β-lactamases, although the identities were relatively low (36.90%-41.30%) (Fig. 1B). No sequences in our dataset shared similarity with Class B, C, or D β-lactamases.

**Figure 1.**
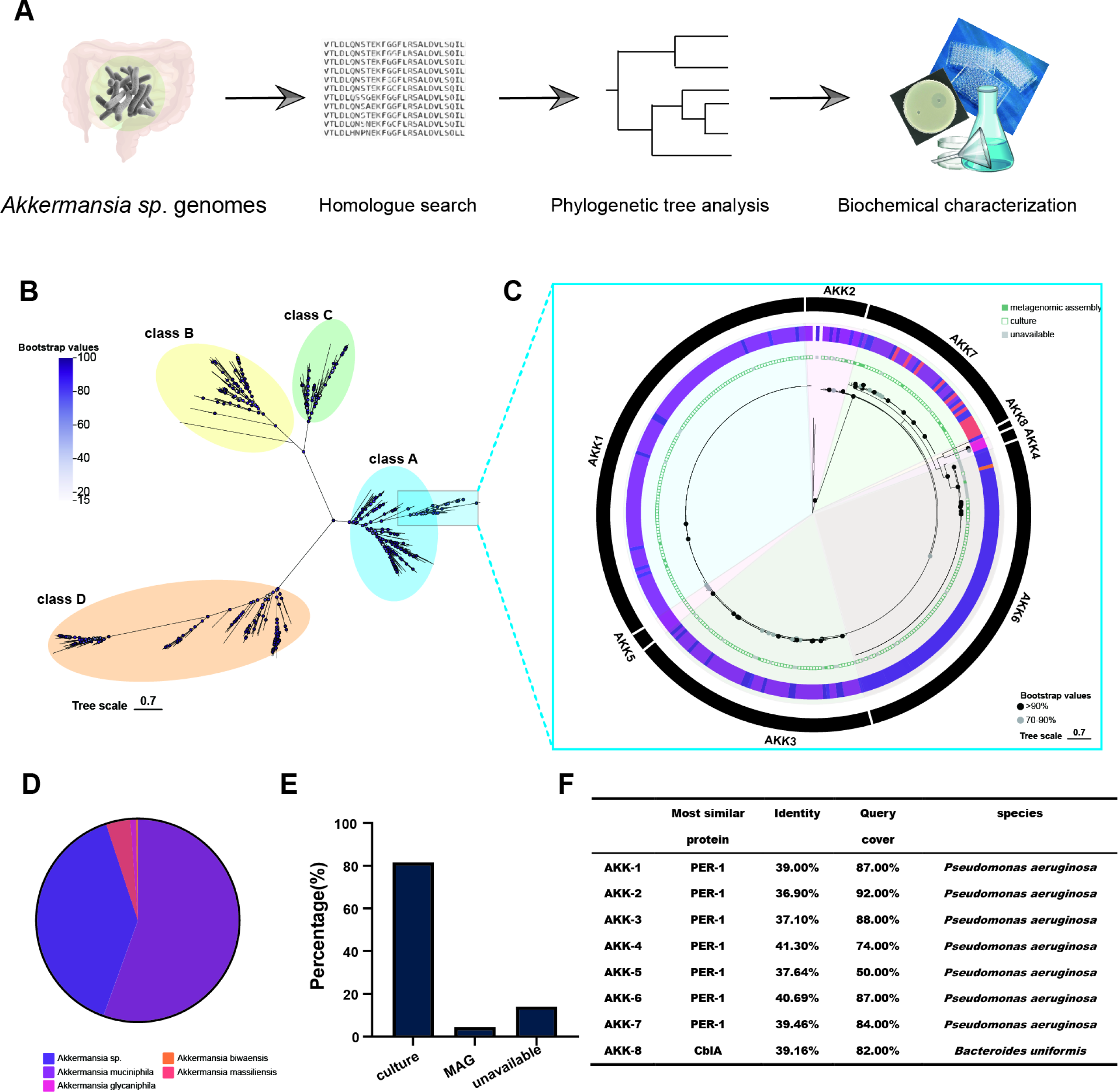
(A) The flow chart of β-lactamase mining process from *Akkermansia* species. (B) Phylogenetic tree analysis of β-lactamase from *Akkermansia* species and β-lactamases from β-lactamase Database (BLDB). Representative sequences were selected using CD-HIT at 80% sequence identity (100 bootstrap replications). (C) The phylogenetic tree analysis of β-lactamase from *Akkermansia* species. 335 predicted β-lactamase from *Akkermansia* species were used for analysis. The innermost circle represents the source of the genome containing the protein (metagenome/pure culture). The middle circle represents the species to which the protein belongs. The outermost circle represents the position of the protein represented by the representative sequence (AKK1/2/3/4/5/6/7/8) on the phylogenetic tree. (D) Pie chart showing the species distribution of 335 predicted β-lactamase. (E) Percentage of predicted β-lactamase sources (culturable or MAG). (F) The information of β-lactamase from BLDB which share highest sequence identity with selected eight representative β-lactamase from *Akkermansia* species.

To further analyze the evolutionary relationship between predicted β-lactamases from *Akkermansia* sp. and BLDB, phylogenetic tree analysis was conducted. We used CD-hit (cut-off value = 0.8) to select representative β-lactamases for phylogenetic tree analysis (Fig. 1C). Notably, the 335 matched proteins from *Akkermansia* species were classified into 8 separate clusters, represented by 8 proteins (AKK-1 to AKK-8). The phylogenetic tree showed that β-lactamases formed 4 distinct clades corresponding to the Class A, B, C, and D families. The 8 proteins formed a small, separate, and distant clade within the Class A clade. They exhibited a closer evolutionary relationship with IBC-2 (*Pseudomonas aeruginosa*), CGA-1 (*Chryseobacterium gleum*), PER-2 (*Citrobacter freundii*), and PER-1 (*Acinetobacter baumannii*) (Fig. 1B). However, the long branches indicated their relatively distant evolutionary relationship with known β-lactamases.

AKK-1, AKK-2, AKK-3, AKK-4, AKK-5, AKK-6, AKK-7, and AKK-8 represented clusters containing 112, 16, 62, 3, 5, 89, 46, and 2 sequences, respectively (Fig. 1C). A detailed phylogenetic tree was constructed using 335 matched proteins from *Akkermansia* species. Proteins from the same cluster classified by CD-hit were also in the same clade in the phylogenetic tree, indicating that the CD-hit clusters correspond to the phylogenetic tree results. Proteins from the AKK-1, AKK-3, and AKK-5 clades were evolutionarily closer, with many members having short branch lengths, indicating their high sequence identity. Most proteins from the eight clusters were from *Akkermansia muciniphila* (55.52%) or *Akkermansia* sp. (25.37%), while AKK-4 and AKK-8 clusters existed in *Akkermansia glycaniphila*, AKK-6 in *Akkermansia biwaensis,* and the AKK-7 cluster in *Akkermansia massiliensis* (Fig. 1C and D). Among these species, the majority are cultivable microbes (81.19%), while 4.78% are uncultivable microbes, and the remaining 14.02% are unidentified microbes (Fig. 1C and E). Interestingly, although metagenome-assembled genomes (MAGs) comprised the highest proportion of the dataset, they contained the fewest β-lactamase sequences. This may be due to insufficient sequencing depth and incomplete genomes in the MAGs.

Our systematic mining found that β-lactamases from *Akkermansia* species shared low similarity and had a distant phylogenetic relationship with known Class A β-lactamases. As far as we know, there are no reports on the biochemical function of β-lactamases from *Akkermansia* species. Considering the risk of resistance gene dissemination is a critical prerequisite for the application of probiotics, especially *Akkermansia* species, we have chosen eight representative β-lactamases from *Akkermansia* species for gene synthesis and protein functional identification (Fig. 1F).

### 2.2 Antibiotic susceptibility of *E. coli* carrying β-lactamases genes from *Akkermansia* sp

The eight β-lactamases from *Akkermansia* species exhibited considerable differences in sequence similarity, signal peptides, and sequence length. They showed diverse sequence similarities with each other: AKK-1 (49.23%-88.73%), AKK-2 (45.09%-92.95%), AKK-3 (48.23%-92.95%), AKK-4 (42.38%-51.89%), AKK-5 (45.82%-99.14%), AKK-6 (49.21%-66.43%), AKK-7 (48.05%-88.73%), and AKK-8 (42.38%-49.23%) (Fig. 2B). Notably, AKK-8 and AKK-4 had the lowest sequence similarity with each other. Despite some sequences showing high sequence similarity, they differed in other aspects. For example, AKK-3 shared 99.14% identity with AKK-5, but AKK-5 had a 244-amino acid uncharacterized region at the N-terminal. AKK-1 shared 92.95% identity with AKK-3, but AKK-3 had an N-terminal signal peptide (Fig. 2A). Notably, AKK-3, AKK-4, AKK-6, and AKK-7 had signal peptides, indicating they are secreted into the periplasmic or extracellular environments (Fig. 2A). This suggests that β-lactamases from *Akkermansia* species might also influence the microenvironments in the host.

**Figure 2.**
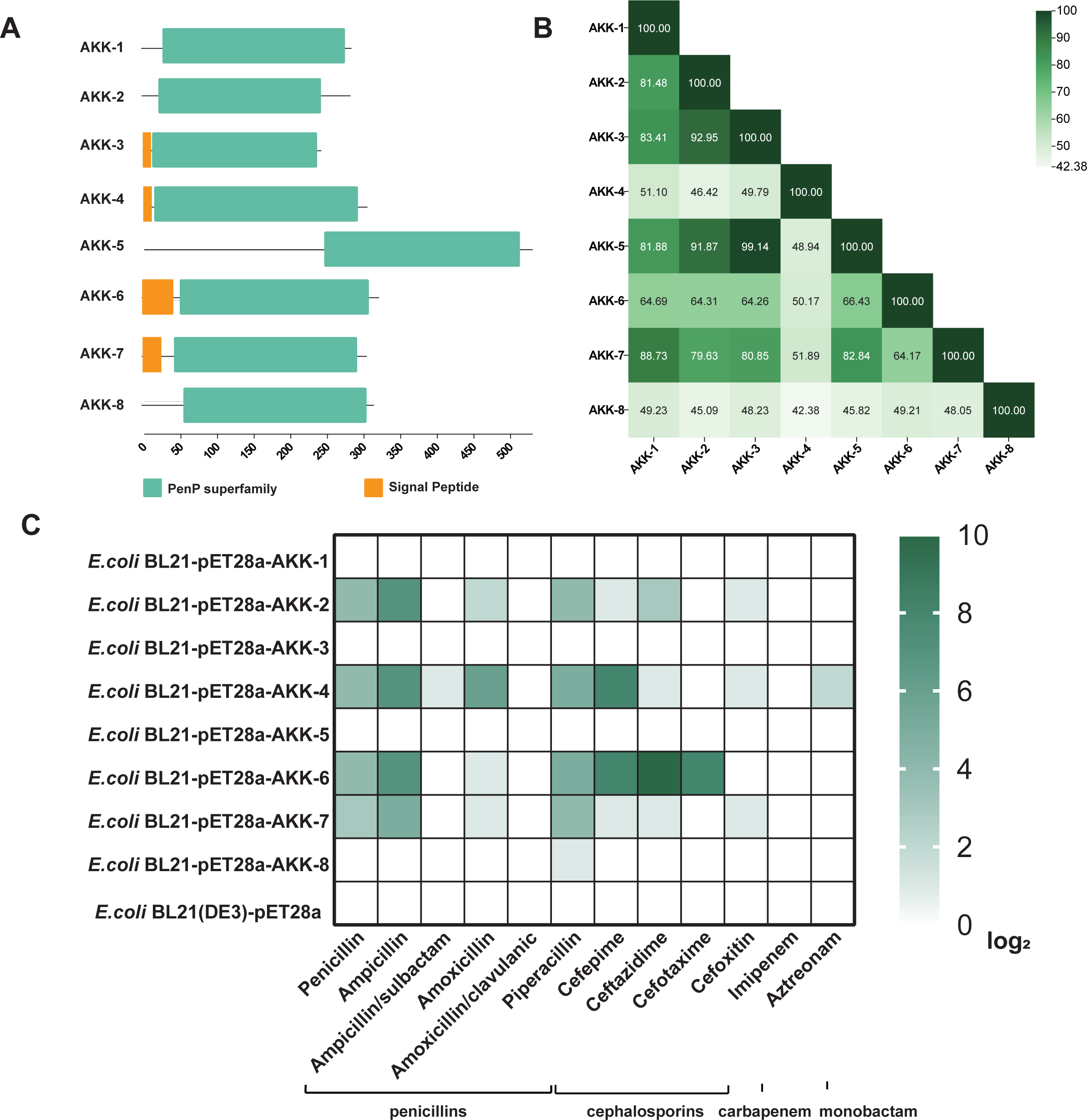
(A) Domain architecture of eight representative β-lactamase from *Akkermansia* species. Orange and green represent signal peptides and PenP superfamily, respectively. (B) A heatmap showing sequence identity between eight representative β-lactamase from *Akkermansia* species. (C) MICs of 10 β-lactam antibiotics and 2 β-lactamase inhibitor combinations (ampicillin/sulbactam and amoxicillin/clavulanic acid) measured for *E. coli* with empty pET28a or eight representative β-lactamase from *Akkermansia* species.

Carrying the β-lactamase from *Akkermansia* species resulted in changes in the minimum inhibitory concentration (MIC) of *E. coli* against β-lactams (penicillins, cephalosporins, and monobactams). *E. coli* with pET28a-AKK-4 showed increased MIC values against β-lactams from the penicillin, cephalosporin, and monobactam classes. *E. coli* with pET28a-AKK-2, pET28a-AKK-6, and pET28a-AKK-7 showed increased MIC values against β-lactams from the penicillin and cephalosporin classes, but not the monobactam class (Fig. 2C). Additionally, *E. coli* with pET28a-AKK-8 only showed an increased MIC value against piperacillin from the penicillin class, but not against other classes. Notably, AKK-6 showed the most significant increase in MIC values for cefepime, ceftazidime, and cefotaxime, with enhancements of 256, 1024, and 256 times compared to the control, respectively (supplementary material 2). Among the eight β-lactamases, AKK-4 exhibited the broadest spectrum of antibiotic resistance, showing resistance changes to eight antibiotics. On the other hand, AKK-8 showed the narrowest spectrum of antibiotic resistance, with resistance changes observed only for piperacillin, with a two-fold increase compared to the control (Fig. 2C and supplementary material 2).

β-lactamase inhibitors reduced resistance to β-lactamase-carrying *E. coli*. For instance, the MIC values of *E. coli* carrying AKK-2, AKK-6 and AKK-7 against sulbactam/ampicillin were reduced to the same level as the control, and AKK-4 MIC decreased by 32-fold (Fig. 2C and supplementary material 2). This indicated that the combined treatment of β-lactam drugs and inhibitors remains effective.

### 2.3 Biochemical characterization of β-lactamases genes from *Akkermansia* sp

To further understand the biochemical properties of β-lactamase from *Akkermansia* sp., we conducted protein expression, purification, and enzymatic characterization. During purification, AKK-1 and AKK-5 were not expressed soluble, while AKK-3 and AKK-8 had insufficient expression, leading to unsuccessful purification. However, four proteins (AKK-2/4/6/7) were successfully purified, showing single bands on SDS-PAGE (Fig. 3A). The molecular weights of AKK-2, AKK-4, AKK-6, and AKK-7 were approximately 31 kDa, 30 kDa, 32 kDa, and 30 kDa, respectively, close to their predicted molecular weights. The purity of the purified proteins exceeded 95%, facilitating further enzymatic characterization studies.

**Figure 3.**
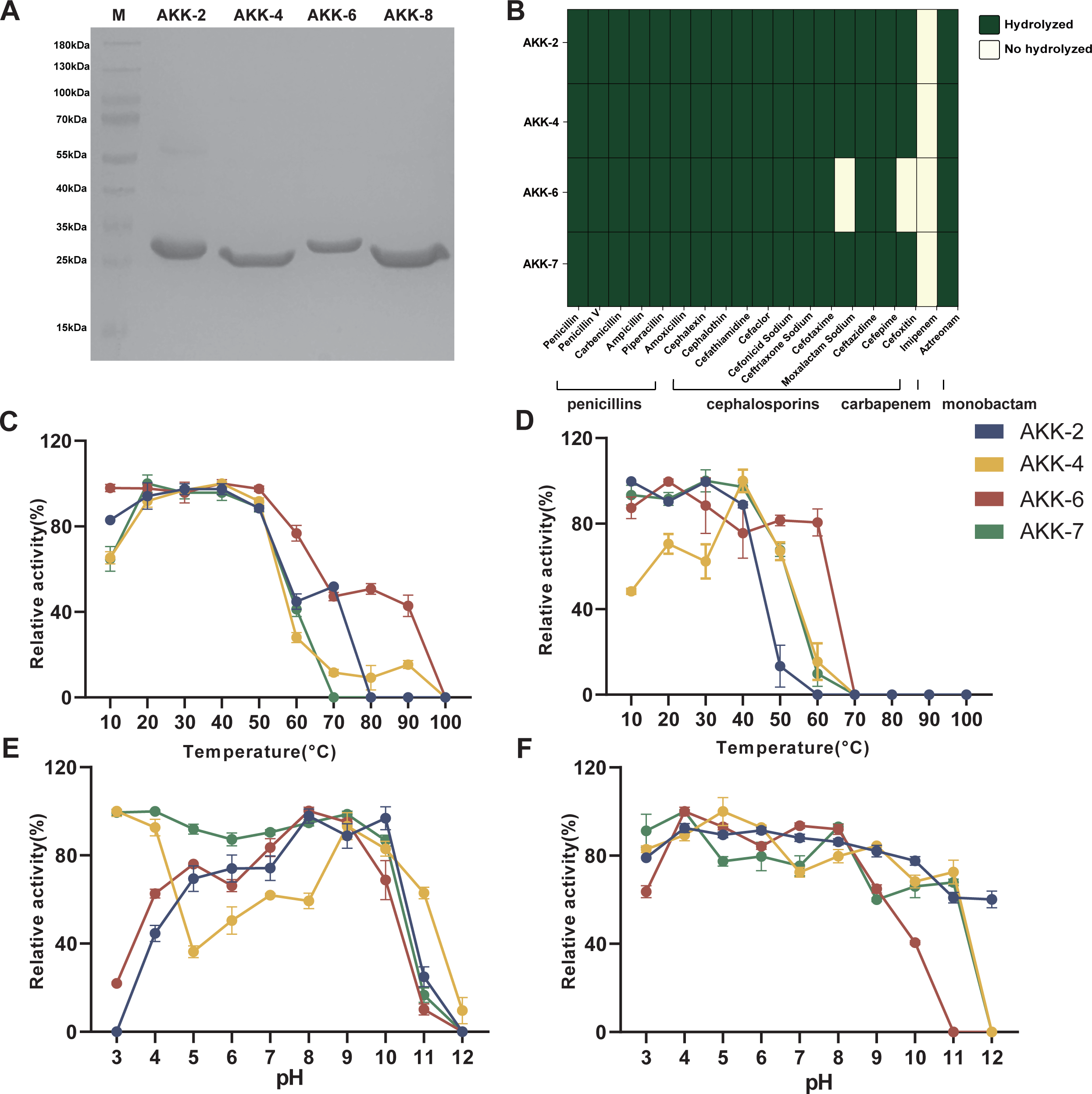
(A) SDS-PAGE analysis of four purified β-lactamase (AKK-2, AKK-4, AKK-6 and AKK-7). (B) Heatmap showing weather β-lactam antibiotics could be hydrolyzed by four purified β-lactamase (AKK-2, AKK-4, AKK-6 and AKK-7). (C) Optimum temperature of four of β-lactamase. (D) Thermostability of the four β-lactamase. (E) Optimum pH for the four β-lactamase. (F) pH stability of the four β-lactamase. Different colors represented different β-lactamase (blue: AKK-2; yellow: AKK-4; red: AKK-6; green: AKK-7).

The optimum reaction temperatures for the four β-lactamases fall within the range of 20-40°C (Fig. 3C): AKK-2 at 30°C, AKK-4 and AKK-6 at 40°C, and AKK-7 at 20°C. Moreover, these proteins retain over 75% activity at temperatures ranging from 10-40°C, with AKK-4 notably stable at 40°C (Fig. 3D). However, its stability decreases at temperatures below this threshold. Regarding pH, AKK-2, AKK-6, and AKK-7 exhibit 80% activity at pH 8-10, 7-9, and 3-10, respectively, while AKK-4 demonstrates higher activity in both alkaline and acidic environments (Fig. 3E). All four proteins exhibit broad pH stability, maintaining over 60% activity at pH 3-9 (Fig. 3F). These optimum reaction temperatures and pH values align well with the intestinal environment where *Akkermansia* sp. is found.

### 2.4 Substrate specificity of β-lactamases genes from *Akkermansia* sp

Substrate specificity of 4 purified β-lactamases were further studied in vitro. All four β-lactamases can degrade three classes of β-lactam antibiotics (penicillins, cephalosporins, and monobactams), but not carbapenems. Notably, AKK-2, AKK-4, and AKK-7 share similar substrate specificity, while AKK-6 cannot degrade moxalactam sodium and cefoxitin (Fig. 3B). β-lactamases from *Akkermansia* exhibit substrate specificity similar to most broad-spectrum Class A β-lactamases, capable of degrading penicillins, cephalosporins, and monobactams. Therefore, AKK-2, AKK-4, and AKK-7 were extended-spectrum β-lactamases.

To further confirm β-lactamase activity, mass spectrometry was utilized to analyze the antibiotics before and after degradation. When antibiotics were treated with the four AKK β-lactamases, inhibition zones of ampicillin, cefotaxime, and aztreonam disappeared, indicating that the antibiotics had lost their activity and were degraded (Fig. 4A, B, D). However, imipenem retained its antimicrobial activity, with the inhibition zone still present (Fig. 4C), indicating that it remained undegraded and active. ESI-MS analysis showed that after enzymatic hydrolysis, signals of ampicillin, cefotaxime, and aztreonam were not detected, and signals corresponding to their hydrolysis products appeared. For example, ampicillin changed from 349 m/z to 367 m/z, cefotaxime from 456 m/z to 473 m/z, and aztreonam from 434 m/z to 452 m/z. The increase in molecular weight by 18 indicates that after hydrolysis of the β-lactam ring, an additional water molecule was added.

**Figure 4.**
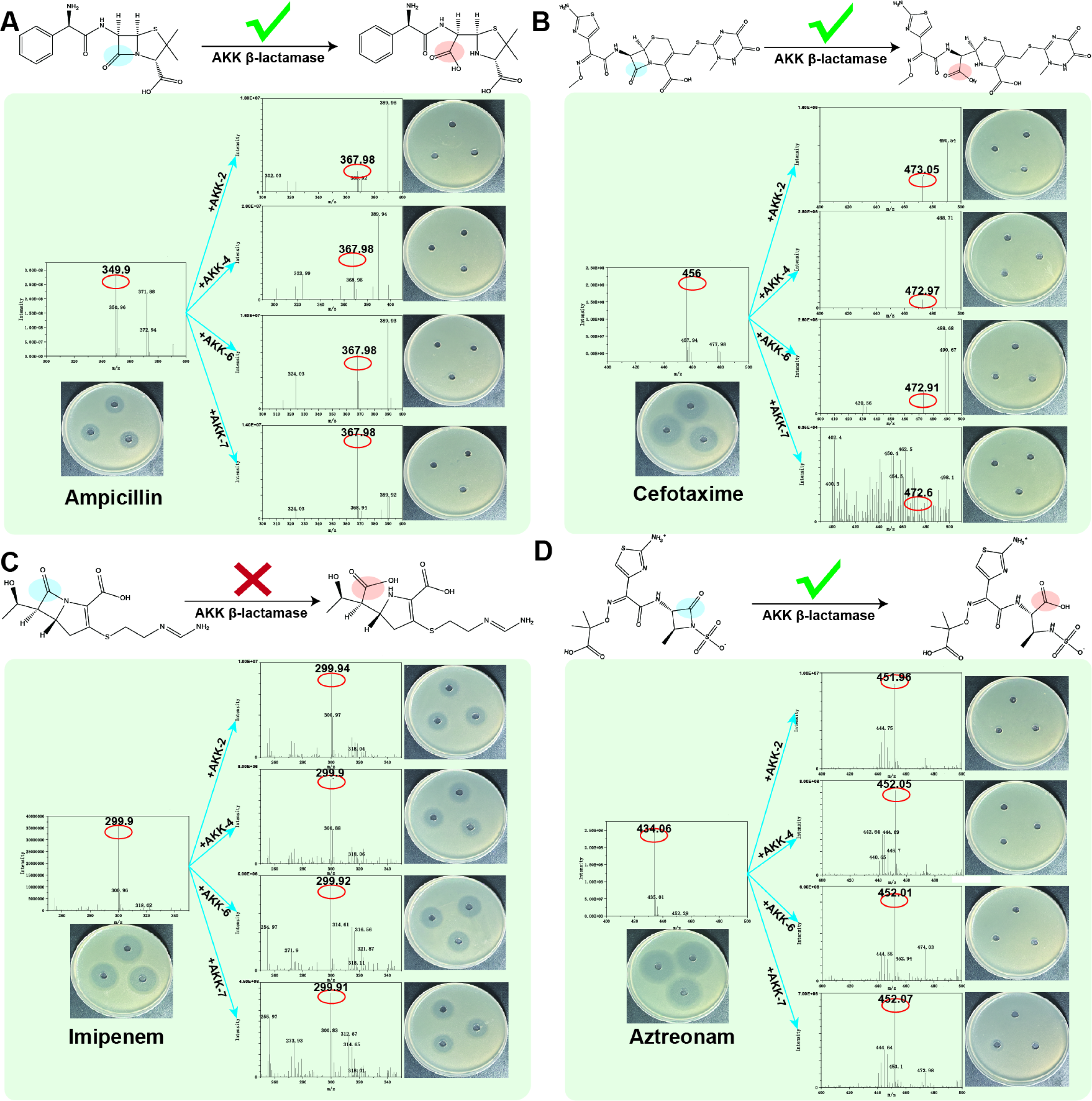
Inhibition zone and ESI-MS analysis of β-lactams before and after degradation by four β-lactamases. (A) ampicillin. (B) cefotaxime. (C) imipenem. (D) aztreonam.

### 2.5 Catalytic amino acid of of β-lactamases from *Akkermansia* sp

The three-dimensional structures of the 8 proteins were predicted using ColabFold, with pLDDT scores ranging from 65 to 95, indicating a high level of confidence (supplementary material 3). Additionally, comparative analysis of the protein 3D structures using the Dali program revealed that the similarity scores among these protein structures are all above 2, indicating that there are highly similar. Binding pockets of the eight β-lactamases were predicted using PRANKweb, and they were also found to be similar to each other (supplementary material 4). Therefore, although the sequence similarity of the eight proteins was different, their 3D structures and binding pockets were similar.

Molecular docking between predicted β-lactamases and ampicillin showed reliable docking results (free energies ranging from -204.441 to -99.2578 kcal/mol) (Fig. 5A). Most β-lactamases interact with substrates primarily through hydrogen bonds, van der Waals forces, and salt bridges (Fig. 5A and supplementary material 5). Considering the low similarity of β-lactamase from *Akkermansia* to known sequences, we further verified its catalytic mechanism. Reported proteins from the Class A family typically utilize serine for catalysis, which is located within the catalytic pocket. Using PRANKweb, we predicted the binding pockets of the eight proteins. Within the pocket, serine appeared only at positions 51, 114, and 221, with conservations of 100%, 50%, and 65%, respectively (Fig. 5B). Among them, the distance between S51 and the substrate (ampicillin) was 2.628 Å. Therefore, we inferred that this amino acid is the catalytic residue.

**Figure 5.**
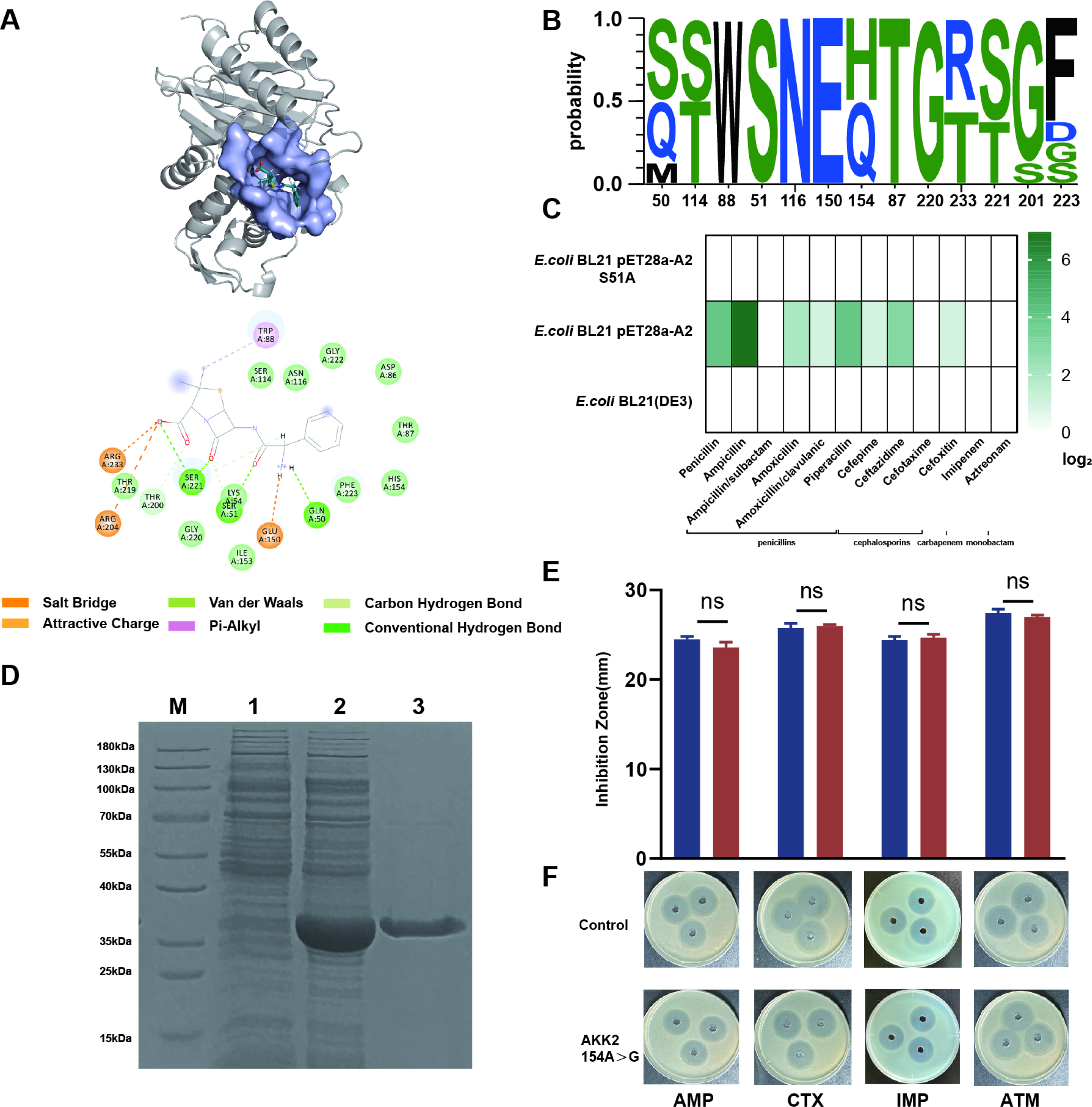
(A) Visualization of the 3D structure of the ampicillin and AKK-2 binding pocket using PyMOL software, and 2D structure of the binding pocket of ampicillin and AKK2. (B) Sequence logos showing sequence conservation of amino acid sequences in binding pocket, The horizontal axis indicates the sequential position of the amino acid. The vertical axis indicates the frequency of occurrence. Each letter represents an amino acid. (C) MICs of 10 β-lactam antibiotics and 2 β-lactamase inhibitor combinations (ampicillin/sulbactam and amoxicillin/clavulanic acid) measured for *E. coli* with empty pET28a or mutated AKK-2(S51A). (D) SDS-PAGE analysis of the purified AKK-2 (S51A). (E) Analysis of the inhibition zone size of the products of 4 β-lactams (AMP: ampicillin; CTX: cefotaxime; IMP: imipenem; ATM: aztreonam) before and after AKK-2 (S51A) hydrolysis. (F) Plate inhibition zone assay showing the antimicrobial activity of 4 β-lactam antibiotics against before and after AKK-2(S51A) hydrolysis.

Through site-directed mutagenesis, we mutated the predicted catalytic serine residue at position 51 in AKK-2. The bacterial resistance phenotype and sensitivity to β-lactam antibiotics remained unchanged in bacteria harboring the mutated gene, similar to *E. coli* BL21 (Fig. 5C). Furthermore, the mutated protein was heterologously expressed and purified using IMAC. The molecular weight of the purified protein matched the predicted size (Fig. 5D). When the purified protein was used to hydrolyze β-lactam antibiotics and subjected to inhibition zone assays, it was observed that the inhibition zones did not disappear (Fig. 5E). The mutated enzyme failed to hydrolyze the antibiotics, and the mixture retained antibacterial activity, indicating that the amino acid is indeed the catalytic residue.

### 2.6 Genomic context of β-lactamases genes from *Akkermansia* sp

After confirming the activity of β-lactamase from *Akkermansia*, a more intriguing question arises: does horizontal gene transfer occur in AKK β-lactamases? Mobile elements or other antibiotic resistance genes often co-occur in the same genomic loci, so we analyzed the occurrence frequencies of protein families in the genomic context. Protein families including the DUF456 domain, DUF3784 domain, tRNA-synt_1 domain, and Peptidase_A8 domain showed occurrence frequencies of 95.31%, 91.55%, 85.92%, and 87.32%, respectively (Fig. 6A). Noteworthy, there were 13 domains with frequencies exceeding 50% in the upstream and downstream environments of AKK proteins, suggesting conserved members around β-lactamase genes (Fig. 6B). Major proteins surrounding the β-lactamase gene include the DUF456 protein (presumed membrane protein), the N-acetyltransferase function protein Acetyltransf_11, and the Haloacid dehalogenase-like hydrolase protein HAD_2. Among these, DUF456 is positioned at -1, opposite the direction of the β-lactamase gene, while Acetyltransf_11 and HAD_2 are positioned downstream, first and second, respectively, in the same direction as the β-lactamase gene (Fig. 6D). However, we did not find any proteins related to drug resistance genes or gene transfer in the genomic context.

**Figure 6.**
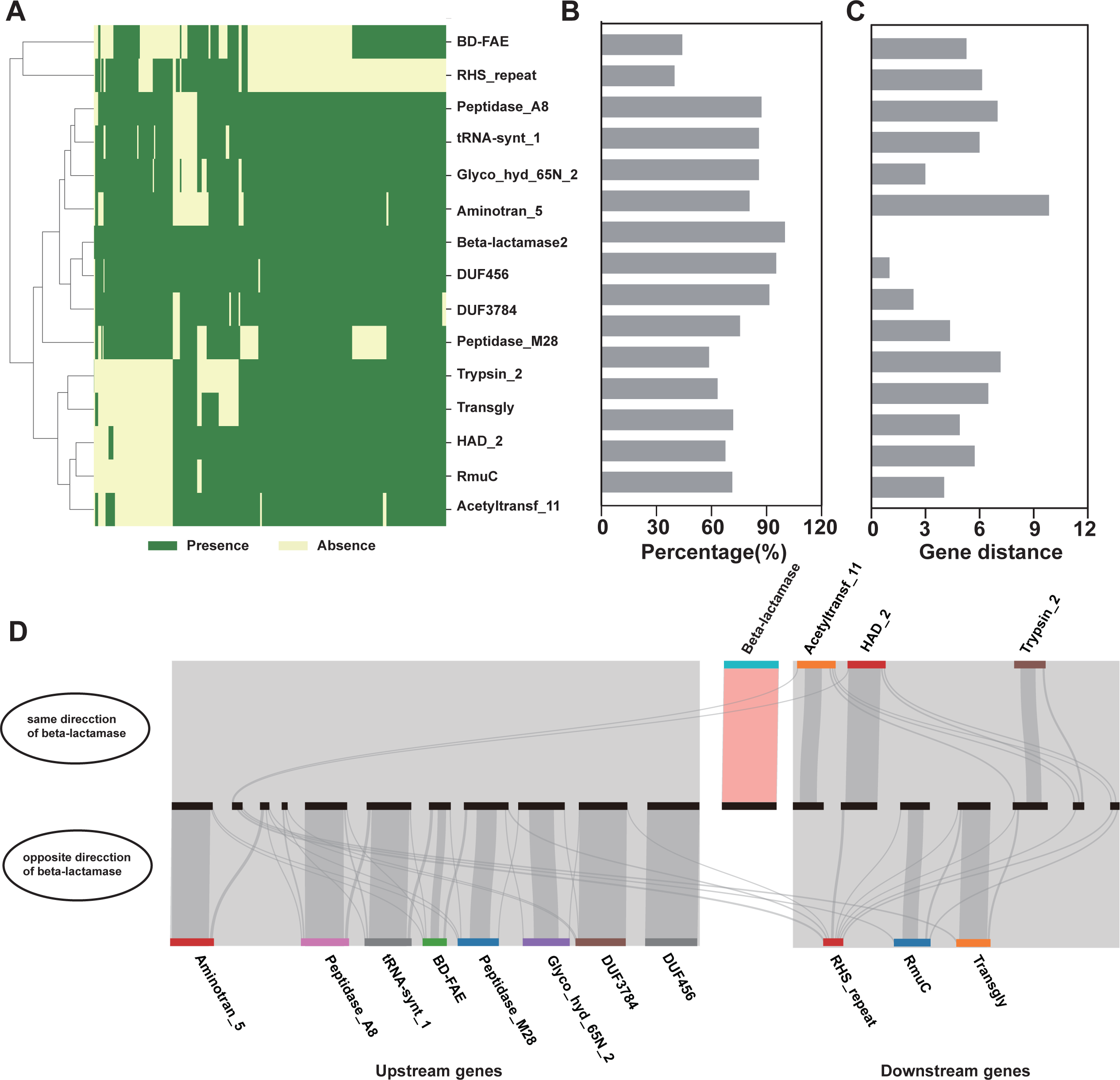
(A) Heatmap analysis representing the distribution of pfam families within the 10 genes upstream and downstream of the β-lactamase gene from *Akkermansia* species. Green indicates the presence of the Pfam family, while light yellow indicates its absence. (B) Percentage distribution of Pfam families within the 10 genes upstream and downstream of the β-lactamase gene. (C) Gene distance between genes from different Pfam families and β-lactamase. (D) Frequency diagram showing the occurrence of various Pfam families in the genomic regions surrounding the β-lactamase gene. Pfam families related to antibiotic resistance are highlighted in pink.

We also compared representative genomic background of 8 β-lactamases across different *Akkermansia* species, including *Akkermansia muciniphila*, *Akkermansia massiliensis*, *Akkermansia biwaensis*, *Akkermansia glycaniphila*, and an unknown *Akkermansia* sp (Fig. 7). Firstly, we found that the genetic backgrounds of β-lactamases are similar among different countries, such as AKK-1 and AKK-5 from the USA and South Korea, and AKK-6, AKK-7, and AKK-8 from Japan, China, the USA, and Australia. Secondly, the genetic backgrounds among different *Akkermansia* species also exhibit similarities, for instance, between *Akkermansia biwaensis* and *Akkermansia massiliensis*, and between *Akkermansia muciniphila* and *Akkermansia massiliensis*. Lastly, similarities are observed in the genetic background across different host sources, including mice, humans, and phascolarctos cinereus. However, we also observed two samples with dissimilar genetic backgrounds, possibly due to limited related research and a lack of similarly recorded genetic backgrounds. Upon observing the gene arrangement upstream and downstream of β-lactamases, we noted significant occurrences of gene rearrangement, loss, and insertion.

**Figure 7.**
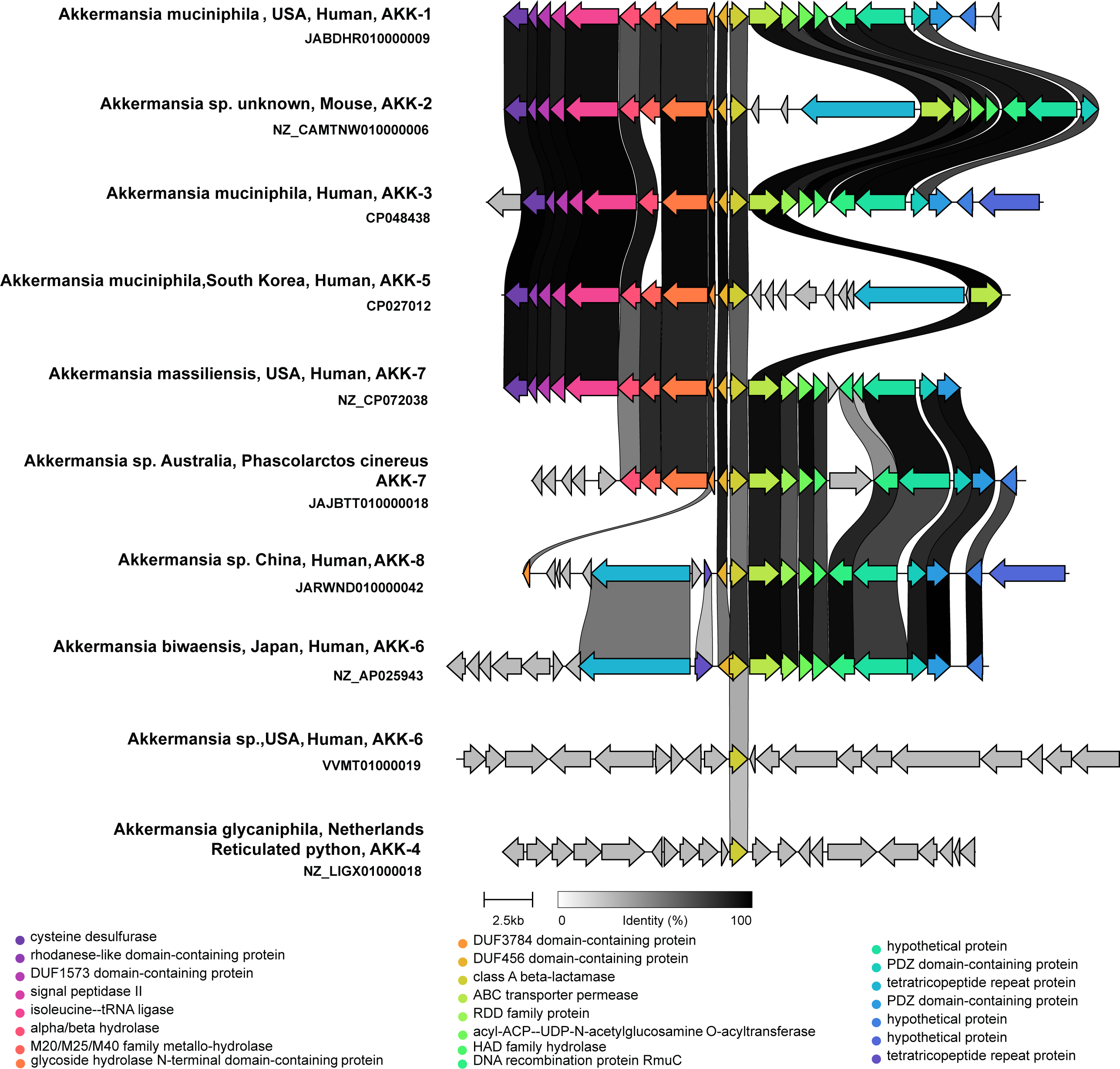
Genomic context linkage analysis of β-lactamase gene loci in representative *Akkermansia* species. Yellow-green arrows represent the β-lactamase gene from *Akkermansia* species. If two sequences from two loci share sequence similarity, they are linked with shadows that change color according to sequence similarity.

## 3. Discussion

### 3.1 Finding of β-lactamases genes from *Akkermansia* sp

*Akkermansia* species is a crucial gut probiotic derived from the *Verrucomicrobia* phylum. The presence of antibiotic resistance genes and their potential for transmission are critical considerations in assessing its safety. This study employed genomic mining techniques to identify eight novel extended-spectrum β-lactamases from *Akkermansia* species sources, sharing similarities ranging from 37.1% to 41.3% with previously identified β-lactamases. This is the first biochemical report of β-lactamases not only in *Akkermansia* species but also in *Verrucomicrobia* phylum. These enzymes confer resistance in *Escherichia coli* to β-lactam antibiotics (ampicillin, benzylpenicillin, and cephalosporins), suggesting that Gram-negative AKK carrying these β-lactamase genes may exhibit resistance accordingly. This hypothesis is consistent with existing reports; for example, Ritesh Kumar et al. identified *Akkermansia* sp. DSM 33459 as resistant to ciprofloxacin, ampicillin, and benzylpenicillin(18). Additionally, studies indicate that A*. muciniphila* DSM 22959 exhibits varying degrees of resistance to antibiotics such as gentamicin, kanamycin, streptomycin, vancomycin, metronidazole, and ciprofloxacin(12, 19, 20). We further validated the minimum inhibitory concentration (MIC) of *A. muciniphila* DSM 22959, which carried AKK-3 and AKK-5, showing resistance to cefotaxime and aztreonam (see supplementary material 6). The presence of β-lactamase genes may assist *Akkermansia* species in surviving β-lactam use, as observed in previous studies, in which increased *Akkermansia* species population increased after use of antibiotics. However, further experimental validation, such as gene knockouts and knock-ins, is necessary to establish the relationship between β-lactamase genes and resistance phenotypes.

### 3.2 Spread of β-lactamases genes from *Akkermansia* sp

Another critical aspect in assessing the safety of *Akkermansia* species is the potential for horizontal gene transfer (HGT) of their carried resistance genes. Currently, we have not found evidence of HGT of β-lactamase genes from *Akkermansia* species to other species, especially pathogenic bacteria. Several observations support this conclusion: (1) No homologs of β-lactamases from *Akkermansia* species (>50% identity) were found in other genera or phyla, indicating that recent horizontal transfer is unlikely. (2) Genomic context analysis revealed the absence of mobile genetic elements (such as transposases, integrases, or insertion sequences) near the β-lactamase genes in *Akkermansia* species, suggesting a low probability of HGT. (3) No antimicrobial resistance genes were found in the genomic context of *Akkermansia* β-lactamases, which typically form gene clusters for coordinated transcription and regulation to respond to antibiotic stress. However, we also found evidence that the GC content of β-lactamase genes is significantly different from that of *Akkermansia* genomes, suggesting a potential horizontal gene transfer event. Therefore, while most evidence suggests that β-lactamase genes from *Akkermansia* species are unlikely to undergo HGT, caution is still warranted, necessitating further safety assessments before their application.

## 4. Materials and methods

### 4.1 Screening for β-lactamases in *Akkermansia* sp

Bacterial genome data of *Akkermansia* genus were retrieved from the NCBI dataset (frozen in June 2023), and the protein sequences were extracted into a single FASTA file for further analysis. Β-lactamase sequences were obtained from the BLDB (http://bldb.eu/) website, and only those with references were collected. The Diamond software was employed to search for β-lactamase homologs in the *Akkermansia* genomes, utilizing the collected β-lactamase sequences as query sequences. The e-value threshold for the Diamond search engine was set to 1e^-10^.

### 4.2 Protein expression and purification

The identified β-lactamase genes were synthesized and subsequently cloned into the pET28a vector, which were transformed into *Escherichia coli* BL21 (DE3) cells. Each recombinant β-lactamase protein had a C-terminal 6xHis-tag sequence for subsequent experiments. *E. coli* BL21 (DE3) cells were cultured overnight, reinoculated (1:1000) with kanamycin, and incubated at 37°C to an OD_600nm_ of 0.6-0.8. Protein expression was induced with 1 mM Isopropyl β-D-Thiogalactoside (IPTG) at 25°C. Cells were then centrifuged, sonicated, and the supernatant purified using immobilized metal affinity chromatography (IMAC) with an imidazole gradient. β-lactamase purity was confirmed by SDS-PAGE.

### 4.3 Minimum inhibitory concentration analysis

The minimum inhibitory concentrations (MICs) of *E. coli* BL21 (DE3) harboring recombinant plasmids were examined using the broth microdilution method for 10 β-lactam antibiotics (penicillin, ampicillin, amoxicillin, piperacillin, cefepime, ceftazidime, cefotaxime, cefoxitin, aztreonam and imipenem) alone or in combination with β-lactamase inhibitors (ampicillin/sulbactam and amoxicillin/clavulanic acid). Overnight cultures of *E. coli* were inoculated into Mueller-Hinton (MH) broth at a final inoculum of 5 × 10^5^ CFU/mL and supplemented with twofold serial dilutions of the antibiotics. After 24 h of incubation at 37°C, the absorbance at OD_600nm_ was measured, and the MIC was determined as the lowest concentration of the antibiotic that inhibited bacterial growth.

### 4.4 Enzyme assay

The enzyme activity of β-lactamase was detected using ampicillin as the substrate. 50 μL of the purified protein was added to 950 μL 120 μg/mL ampicillin solution in 20 mM Tris-HCl buffer (pH 7.0). The solution was thoroughly mixed and incubated at 37°C for 10 min. The reaction was terminated by boiling the reaction solution. The optical density at 235 nm (OD_235nm_) was measured using a spectrophotometer to determine the enzyme activity(^21^). Enzyme activity was defined as the amount of enzyme required to produce 1 μM of penicilloic acid per minute under standard conditions (22). All experiments were performed in triplicate, unless otherwise specified.

### 4.5 Phylogenetic tree analysis of *Akkermansia* β-lactamase

After obtaining the matched β-lactamase candidates from the Diamond analysis in section 2.1, the sequences were clustered using CD-HIT at 80% sequence identity, and the representative sequences were used to build a phylogenetic tree along with the collected β-lactamase sequences using IQ-tree software. Subsequently, the phylogenetic tree was visualized and rendered using the Chiplot program (^23^).

### 4.6 Biochemical characterization of β-lactamase

To investigate the effect of temperature on enzyme activity, the purified β-lactamase was added to ampicillin in 20 mM Tris-HCl buffer (pH 7.0), and the reaction was carried out at 10, 20, 30, 40, 50, 60, 70, 80, 90, and 100°C for 10 min, respectively (24). Subsequently, the enzyme activity was measured according to the method outlined in Section 2.4. To evaluate thermal stability, the purified β-lactamase was incubated in a water bath at 10, 20, 30, 40, 50, 60, 70, 80, 90, and 100°C for 1 h, respectively, after which the enzyme activity was measured.

Ampicillin solutions were prepared using Britton-Robinson pH buffers (pH 3-11). Then, the enzyme activity was measured by the method described in Section 2.4. To test pH stability, the purified β-lactamase was added to different pH buffers and incubated at 4°C for 1 h. Subsequently, the enzyme activity was evaluated.(24).

### 4.7 Substrate specificity of β-lactamase

To evaluate the specificity of the β-lactamases against different β-lactam antibiotics, a total of 19 β-lactam antibiotics were tested. These included 6 penicillins (penicillin, penicillin V, carbenicillin, ampicillin, piperacillin and amoxicillin), 11 cephalosporins (cephalexin, cephalothin, cefathiamidine, cefaclor, cefonicid sodium, ceftriaxone sodium, cefotaxime, moxalactam sodium, ceftazidime, cefepime, and cefoxitin), 1 carbapenem (imipenem), and 1 monobactam (aztreonam). Enzyme activity assay was similar to that described in Section 2.4. Optical density was measured at different wavelengths appropriate for each antibiotic(21), and the antibiotic concentration was calculated using its corresponding molar absorptivity.

### 4.8 Inhibition zone analysis of enzymatic hydrolysis products

The antibacterial activity of 4 β-lactams (ampicillin, cefotaxime, imipenem, and aztreonam) before and after enzymatic hydrolysis was analyzed using the agar diffusion method. 20 μL of *E. coli* were inoculated into LB agar medium containing 1.5%, thoroughly mixed, and poured onto Petri dishes. When the agar solidified, wells of 4.5 mm diameter were created using a hole puncher. The purified β-lactamase was incubated at 37°C with each antibiotic at a concentration of 20 μg/mL for 1 h. Subsequently, the reaction mixture was incorporated into LB agar medium. After incubating the culture dishes at 37°C for 16 hours, the hydrolysis zones were observed.

### 4.9 ESI-MS analysis of enzymatic hydrolysis products

Electrospray ionization mass spectrometry (ESI-MS) was utilized to verify whether the purified β-lactamase hydrolyzed β-lactam antibiotics (ampicillin, cefotaxime, imipenem, and aztreonam). 50 μL of the purified enzyme was added to 950 μL 50 μg/mL β-lactam antibiotic solution, and the reaction mixture was incubated at 37°C for 30 min, followed by filtration through a 0.22 μm micromembrane filter. The mass spectrometry scan range was set at 100 to 1000 m/z, with a curtain gas pressure of 40 psi, and ion source gas 1 and gas 2 were set to 30 psi each. The electrospray ionization (ESI) voltage was set to 5500 V for positive mode (ESI+) and 4500 V for negative mode (ESI-), and the drying gas temperature and flow rate were maintained at 450°C and 5 L·min-1, respectively.

### 4.10 3D structure prediction and molecular docking

The 3D structure of the β-lactamase was predicted using Colabfold with the following parameter settings: number_recycles 50, max_msa 512-1024, number_relax 5, rank_plddt, and number_models 5(^25^). The protein structures were visualized using PyMOL(^26^) . The binding pocket on the β-lactamase was predicted using the prankweb server (https://prankweb.cz/). Molecular docking between the β-lactamase and β-lactam antibiotics (ampicillin, cefotaxime, imipenem, and aztreonam) was carried out using Libdock, and the docking result with the lowest Gibbs free energy was selected for subsequent analysis(^27, 28^).

### 4.11 Site-directed mutagenesis

To further investigate the catalytic mechanism of the β-lactamase, site-directed mutagenesis was performed. The mutant plasmids were transformed into *E. coli* BL21 (DE3), and the mutations were confirmed by sequencing. The procedures for plasmid construction, protein expression and purification, enzyme activity determination, and minimum inhibitory concentration (MIC) determination were consistent with those described in Sections 2.2, 2.3, and 2.4, respectively. MIC and inhibition zone analysis of *E. coli* BL21 (DE3) with mutants were conducted (section 2.3 and section 2.8).

### 4.12 Genomic context of *Akkermansia* β-lactamase

To study the genomic context of β-lactamase from *Akkermansia* sp., the genomic information of β-lactamase from *Akkermansia* sp. was downloaded from the NCBI database. Ten protein sequences upstream and downstream of the β-lactamase gene were extracted from each individual genome. The hmmer software was then used to identify the Pfam protein family of each extracted protein sequence (E-value: 1e^-10^, minimum matched length: 50). The occurrences of Pfam families in the genomes were summarized, retaining only those with an occurrence rate greater than 10%. Finally, a heatmap was utilized to visually represent the presence of Pfam protein families within the genomic context of the β-lactamase gene. The linkages between various representations of these analyses were visualized using the clinker online website (https://cagecat.bioinformatics.nl/).

## Acknowledgement

This work was founded by Sichuan key research and development program (2023YFS0384), the Open Project Program of Irradiation Preservation Key Laboratory of Sichuan Province, Sichuan Institute of Atomic Energy (NO. FZBC2022003), Zunyi Technology and Big data Bureau, Moutai institute Joint Science and Technology Research and Development Project (ZSKHHZ [2021] No.320), open project of Anti-infective Agent Creation Engineering Research Centre of Sichuan Province (AAC2023010 and AAC2023013), Research Foundation for Scientific Scholars of Moutai Institute (mygccrc[2022] No.066) .This study was supported by the grants from National Natural Science Foundation of China (82372290 and 31801143), Natural Science Foundation of Sichuan Province (2023NSFSC1467, 2023NSFSC1237) and Enzyme Resources Sharing and Service Platform of Sichuan Province.

